# Uterus and cervix anatomical changes and cervix stiffness evolution throughout pregnancy

**DOI:** 10.1101/2024.05.01.592023

**Authors:** Erin M. Louwagie, Serena R. Russell, Jacqueline C. Hairston, Claire Nottman, Chia-Ling Nhan-Chang, Karin Fuchs, Cynthia Gyamfi-Bannerman, Whitney Booker, Maria Andrikopoulou, Alexander Friedman, Noelia Zork, Ronald Wapner, Joy Vink, Mirella Mourad, Helen M. Feltovich, Michael D. House, Kristin M. Myers

## Abstract

The coordinated biomechanical performance, such as uterine stretch and cervical barrier function, within maternal reproductive tissues facilitates healthy human pregnancy and birth. Quantifying normal biomechanical function and detecting potentially detrimental biomechanical dysfunction (e.g., cervical insufficiency, uterine overdistention, premature rupture of membranes) is difficult, largely due to minimal data on the shape and size of maternal anatomy and material properties of tissue across gestation. This study quantitates key structural features of human pregnancy to fill this knowledge gap and facilitate three-dimensional modeling for biomechanical pregnancy simulations to deeply explore pregnancy and childbirth. These measurements include the longitudinal assessment of uterine and cervical dimensions, fetal weight, and cervical stiffness in 47 low-risk pregnancies at four time points during gestation (late first, middle second, late second, and middle third trimesters). The uterine and cervical size were measured via 2-dimensional ultrasound, and cervical stiffness was measured via cervical aspiration. Trends in uterine and cervical measurements were assessed as time-course slopes across pregnancy and between gestational time points, accounting for specific participants. Patient-specific computational solid models of the uterus and cervix, generated from the ultrasonic measurements, were used to estimate deformed uterocervical volume. Results show that for this low-risk cohort, the uterus grows fastest in the inferior-superior direction from the late first to middle second trimester and fastest in the anterior-posterior and left-right direction between the middle and late second trimester. Contemporaneously, the cervix softens and shortens. It softens fastest from the late first to the middle second trimester and shortens fastest between the late second and middle third trimester. Alongside the fetal weight estimated from ultrasonic measurements, this work presents holistic maternal and fetal patient-specific biomechanical measurements across gestation.

## Introduction

Pregnancy and parturition are remarkably mechanical processes, orchestrated by the growth, remodeling, and activity of maternal reproductive and gestational tissues. From the nonpregnant state to term gestation, the uterus, a thick-walled organ comprised predominantly of smooth muscle cells, sheathed in extracellular matrix (ECM), must remain quiescent as it grows in mass from about 70 to 1100 grams and stretches to accommodate intrauterine contents of around 10 milliliters in the nonpregnant state to 5 liters [1, 2]. As the uterus is stretched, the cervix, a collagen-rich organ connecting the intrauterine cavity to the vaginal canal, must remain closed to retain the fetus as it develops [3]. Similarly, the fetal membranes, a thin multi-layer tissue enclosing the fetus and amniotic fluid, must remain intact to provide mechanical support and protection from infection to the growing fetus [4]. At the time of parturition, these functions must all be reversed: the fetal membranes rupture, the cervix dilates, and the uterus contracts to allow for vaginal delivery. Though the exact mechanism(s) controlling these changes is unknown, it is clear that the mechanical failure and mistiming of these tissues can have catastrophic consequences.

In the prenatal period, one of the most common and dangerous outcomes is preterm birth (PTB, live birth before 37 weeks gestation). Globally, more than 1 in 10 babies is born preterm, and PTB-related complications are the leading cause of death for children under 5 years of age [5]. Cited causes of PTB with biomechanical implications are uterine overdistention, preterm premature rupture of membranes, and cervical insufficiency [6]. Still, the pathophysiology and interplay of these conditions remain unknown, and 70-80% of PTB is spontaneous and unexplained [6, 7]. The inability to accurately predict why and in whom PTB will occur has led to a lack of effective therapies for PTB prevention [8]. To truly understand how maternal reproductive anatomy supports the intrauterine load in uncomplicated pregnancies and target therapies in pregnancies at high risk for PTB, we must pursue computational approaches.

Pregnancy is a protected environment. Thus,*in-silico* approaches based on non-invasive measurements, such as ultrasound, are one of the only viable options for elucidating the biomechanical features of the pregnancy environment. Large institutional bodies have recognized the importance of computational modeling and simulation in clinical trials and device design, with the United States Food and Drug Administration (FDA) forming the Modeling and Simulation Working Group in 2016 and the American Society of Mechanical Engineers (ASME) releasing a standard for the verification and validation of computational models specifically for medical devices in 2018 [9, 10]. Advancements in *in-silico* methods for clinical study have facilitated vast research in fields such as cardiovascular, cartilage, and tumor growth biomechanics [11–13]. These methods can be used as digital twins to predict disease progression and tailor medical devices to individual patient needs. In pregnancy, these patient-specific models could include details such as previous cesarean section scars [14, 15]. The key roadblock in using computational methods to study and design devices for pregnancy complications, such as PTB, is a lack of data on biomechanical changes in human maternal reproductive tissues across gestation.

The first *in-vivo* data on maternal reproductive anatomy size and shape during pregnancy was published in 1950, where 15 pregnant uteri were imaged via x-ray at regular intervals from 18 weeks of gestation until term, reporting outlines of the outer uterus and ratios of the inferior-superior to anterior-posterior uterine lengths [16]. However, x-ray imaging does not capture soft tissues well and is not commonly used in prenatal care today [16]. In 2010, ultrasound techniques were used to estimate uterine wall tension across gestation in 320 pregnancies (294 term singletons, 15 preterm singletons, and 11 twins), collecting measurements of inferior-superior, anterior-posterior, and left-right uterine diameter with anterior uterine wall thickness, though no cervical measurements were included [17]. The cervix is the most scrutinized of the maternal anatomic features during pregnancy, as cervical length can be used as a predictive tool for PTB, and time-course studies of cervical length and stiffness have been reported [18, 19]. In 2021, our team published the first datasets on uterine and cervical dimension measurements, cervical stiffness data, and computer-aided design (CAD) models of the uterus and cervix across gestation in 29 uncomplicated pregnancies, making all data and models publicly available to create wider opportunity for *in-silico* pregnancy research [20, 21].

In this study, we build upon previous work to quantify anatomical and biomechanical changes in the uterus and cervix across gestation in pregnant participants at low-risk for preterm birth. Our approach is to parametrically assess the size and shape of the uterus and cervix through a clinically implementable two-dimensional (2D) ultrasound protocol and measure *in-vivo* cervical stiffness via aspiration [20, 22]. Additionally, we use these parametric measurements to generate patient-specific CAD models across gestation. We present the following for pregnancies at low risk for PTB: 1) parametric dimension measurements of the uterus and cervix across gestation, 2) measurements of cervical stiffness via aspiration (Pregnolia AG, Schlieren, Switzerland), 3) the effects of gestational age on maternal reproductive anatomy with comparisons to previously published data [18, 20, 21], 4) patient-specific parametric CAD models of the uterus and cervix to estimate uterocervical tissue volume (Solidworks, Dassault Systémes, Vélizy-Villacoublay, France), and 5) corresponding measurements of fetal size and amniotic fluid level. This work makes available coordinated, patient-specific measurements of maternal reproductive tissue, the fetus, and cervical stiffness across gestation.

## Methods

A prospective, time-course, observational study of human pregnancy in participants at low risk for PTB was conducted. Maternal reproductive anatomy, fetal size, and amniotic fluid level were measured via two-dimensional (2D) ultrasound and cervical stiffness using the Pregnolia system (Pregnolia AG, Schlieren, Switzerland). Measurements were collected at four time points during gestation: late first trimester (L1, 9w3d-15w3d), middle second trimester (M2, 17w3d-20w5d), late second trimester (L2, 23w3d-27w1d), and middle third trimester (M3, 33w4d-38w0d). Linear regressions were fit to the data to find relationships with gestational age, and T-tests were performed to analyze the effect of parity. All findings were compared to existing datasets on normal pregnancy.

### Study Design

This was a prospective observational study of ultrasound dimension and cervical stiffness measurements in participants at low risk for PTB at 9w3d-15w3d, 17w3d-20w5d, 23w3d-27w1d, and 33w4d-38w0d gestation.

### Participants

Participants were recruited from prenatal care patients at a single tertiary care center in New York, New York. Participants were approached after being introduced to the study by their primary obstetrician and consented to all study protocols prior to research participation. Fifty participants aged 23-41 were recruited and participated in the study from April 8, 2019 to June 17, 2023. Inclusion criteria included being 18 years of age or older, carrying an uncomplicated singleton gestation, and being able to provide informed consent. Exclusion criteria included a current IVF pregnancy, multifetal reduction, history of PTB, history of cervical surgery (LEEP, cone biopsy, trachelectomy, cerclage), history of cervical shortening in the current pregnancy, history of significant vaginal bleeding during pregnancy, history of any major abdominal/uterine surgery, history of cesarean section, abnormal pap smear, persistent cramping, persistent uterine contractions, vaginal bleeding at the time of consent, uterine anomalies, systemic or vaginal infections at the time of consent, on progesterone in the first trimester, fibroids, placenta previa or abnormal placentation, ovarian cysts (other than a corpus luteal cyst), and anything in vaginal canal in last 24 hours at the time of the first research visit. This study was approved by the institutional review board at Columbia University Irving Medical Center (CUIMC), and each participant provided written informed consent.

Participant age, height, weight, race, ethnicity, patient history, social history, and obstetric history were recorded for each patient. At each prenatal research visit, participants reported their weight, pregnancy complications since their last visit, and whether they’d had intercourse or other vaginal transition in the past 24 hours (tampon, yeast medication, etc.). After the participant delivered, the gestational age at delivery and mode of delivery (cesarean or vaginal) was recorded. Fifty participants were recruited to and participated in the study. Two participants delivered preterm (before 37 weeks gestation), and one miscarried. These participants were excluded from the analysis. Patient demographics for the remaining 47 participants are reported in table 1.

**Table 1.** Participant demographics and parity. Number of subjects, N, and percentage of total subjects (in parentheses) in each group.

| Demographic |  | N = 47 | (%) |
| --- | --- | --- | --- |
| Age (years): | Mean (± STD) | 32 (± 4) | |
|  | Range | 23-41 |  |
| Parity: | Nulliparous | 30 | (63.8) |
|  | Multiparous | 17 | (36.2) |
| Ethnicity: | Hispanic | 12 | (25.5) |
|  | Non-Hispanic | 33 | (70.2) |
|  | Unknown/Not Reported | 2 | (4.3) |
| Race: | Asian | 3 | (6.4) |
|  | Black/African American | 6 | (12.8) |
|  | White | 33 | (70.2) |
|  | Unknown/Not Reported | 5 | (10.6) |

### Ultrasound and Cervical Stiffness Measurements

Cervical stiffness was measured using the Pregnolia system (Pregnolia AG, Schlieren, Switzerland). This is an aspiration device administered during a speculum exam. Clinicians were trained to use the device following the “instructions for use” documentation and training videos provided by the Pregnolia website [23]. The system is operated via a control unit containing a vacuum pump attached to the probe by flexible tubes. The vacuum is turned on via a foot pedal, creating a negative pressure at the probe head, and the clinician brings the probe head into contact with the distal anterior lip of the cervix. Once contact is established, the clinician slides the probe handle to the middle position, and the negative pressure is increased to pull the cervical tissue 4mm. The vacuum is stopped at this point, and the pressure required to displace the tissue 4mm is recorded. The measurement is performed three times. The average of these values is reported as the aspirated cervical stiffness (aCS).

Sonographers trained on the study protocol performed prenatal ultrasound examinations. Ultrasounds were collected by sonographers (I.K., I.M., and V.P.) trained on the ultrasound acquisition protocol by the material-fetal medicine specialist who initially developed it (C-L.N-C.). Before imaging, participants were asked to empty their bladders. Ultrasoound images were obtained using a GE Voluson E8 (GE Healthcare, Chicago, IL, USA). Standard clinical ultrasonic dimensions of the fetus were measured. Crown-rump length (CRL, cm) was measured at timepoint L1, and estimated fetal weight (EFW, grams) was measured at timepoints M2, L2, and M3. EFW was calculated using the Hadlock I formula, requiring measurements of biparietal diameter (BPD), head circumference (HC), abdominal circumference (AC), and femur length (FL) [24]. The maximum volume pocket (MVP, cm) was collected at all time points, and amniotic fluid index (AFI, cm) was collected at time points L2 and M3, and occasionally at M2. The placenta location was also recorded at all time points.

With the participant in the supine position, the three B-mode research images of maternal anatomy were acquired: transabdominal (TA) sagittal, TA axial, and transvaginal (TV) sagittal. The TA and TV images were collected following the protocol described in our previous work [20]. The TA images were acquired using extended field-of-view ultrasound imaging, where the probe was swept across the abdomen, and adjacent images were automatically registered to produce one long image. The extended field-of-view feature permitted imaging of the full length of the uterus in one image. The sonographer acquiring the images placed calipers on the images to mark the location of the dimension measurements at the time of image acquisition. Precise dimension measurements were later taken by E.M.L. using Fiji ImageJ [25]. Maternal-fetal medicine specialists M.H. and C-L.N-C reviewed images and measurements to verify adequate visualization of the structures under study, including notation of any problematic measurements, such as inaccurate cervical length resulting from a lower uterine segment contraction that distorts the anatomy.

From the TA sagittal ultrasonic image, the longest inferior-superior intrauterine diameter was measured from the fundal to the lower uterine segment endometrium (IS-UD), marking the inferior-superior intrauterine axis (Fig. 1a). Perpendicular to the midpoint of the inferior-superior intrauterine axis, the anterior-posterior intrauterine diameter (AP-UD) was measured (Fig. 1a). The TA axial ultrasonic image provided measurements of the longest left-right intrauterine diameter (LR-UD) (Fig. 1b). The TV sagittal image provided measurements of cervical length (CL), which was measured as the distance between the internal os (where the anterior and posterior cervix meet in the image) and the external os, and the smallest thickness of the lower uterine segment (LUS-UT) (fig. 1c). Additional ultrasonic measurements were collected to generate the patient-specific CAD models, but were not included in statistical analysis (S1 Appendix).

**Fig 1.**
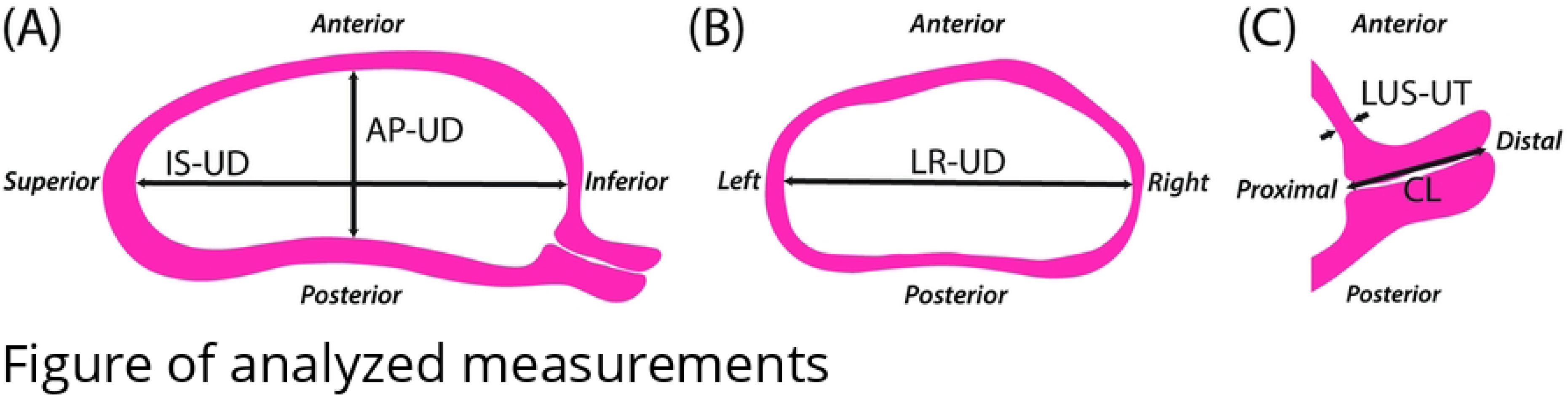
2D ultrasonic measurements of the uterus and cervix. (A) Transabdominal panoramic 2D sagittal ultrasounds measured the inferior-superior (IS-UD) and anterior-posterior (AP-UD) intrauterine diameters. (B) Transabdominal panoramic 2D axial ultrasounds measured the left-right intrauterine diameter (LR-UD). (C) Transvaginal 2D sagittal ultrasounds measured cervical length (CL) and lower uterine segment thickness (LUS-UT). These dimensions are based on previously published measurement protocols [20, 26].

### Estimated Uterocervical Volume

The estimated uterocervical volume (EUV) was found from solid models generated from the maternal ultrasonic dimensions. Parametric patient-specific CAD models were built in Solidworks 2018-19 (Dassault Systémes, Vélizy-Villacoublay, France) using an existing modeling protocol for all participant visits with ultrasonic dimension measurements (fig. 2 [20]. Several updates to the modeling protocol were made, as outlined in S1 Appendix. The EUV was found using the “Mass Properties” tool within Solidworks.

**Fig 2.**
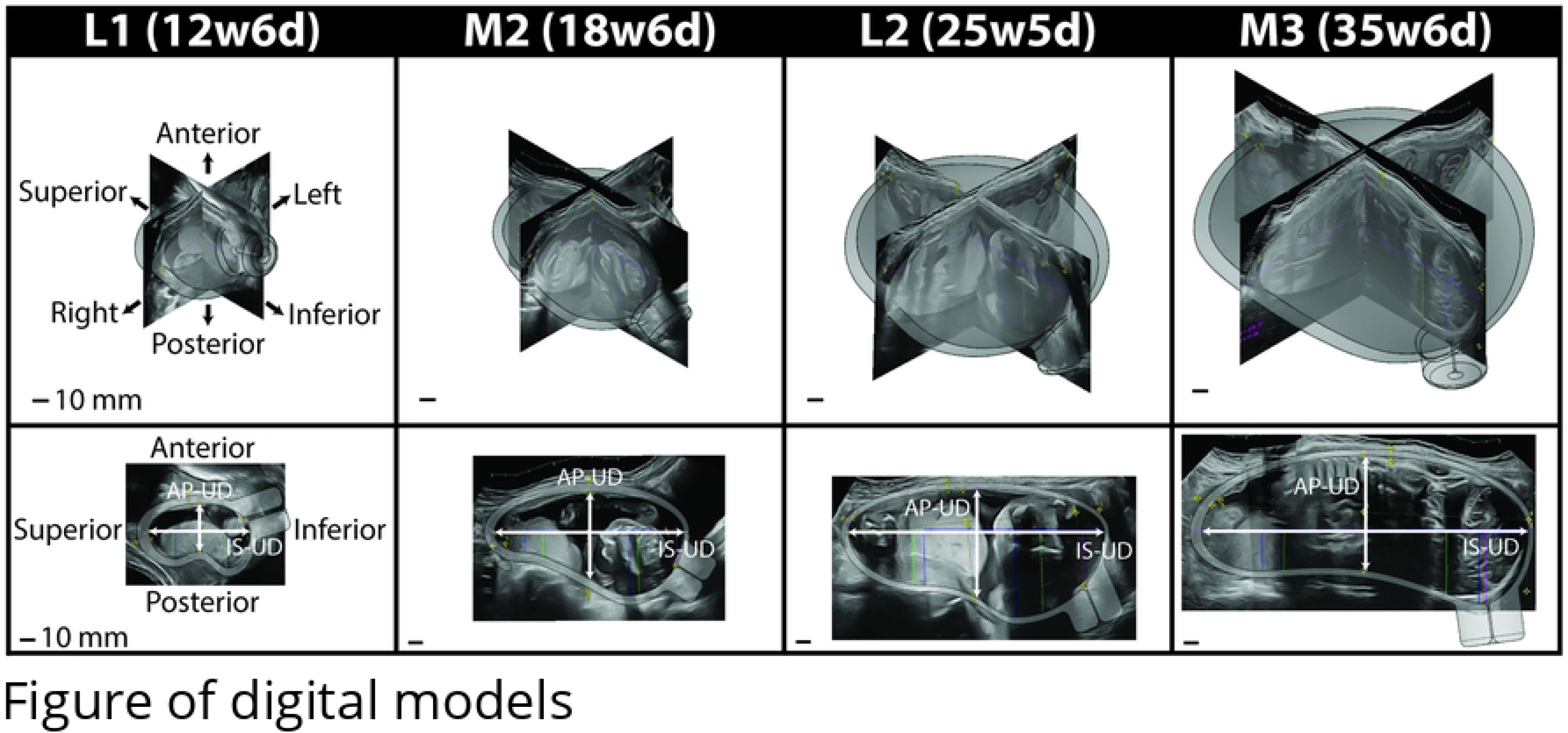
Uterus and cervix CAD models across gestation. Three-dimensional solid CAD models of the uterus and cervix for one participant from the late first to middle third trimester with the corresponding ultrasonic images. The top row is the isometric view, and the bottom is the sagittal view. The inferior-superior (IS-UD) and anterior-posterior (AP-UD) intrauterine diameters are marked in the sagittal view.

### Statistical Analysis

A summary of participant numbers at each visit and those who attended the previous visit is provided in table S1 Table 1. Patient characteristics, uterine dimensions, cervical length, and cervical stiffness were analyzed in RStudio version 1.3.1056 [27]. Ultrasound measurements and cervical stiffness were analyzed using a linear regression accounting for differences in both parity and participant. Welch’s t-tests were used to determine measurement differences between cohorts and visits. For the slope comparisons, the hypothesis is that there is no difference between the measurements from the two time points being assessed. The assessed slopes are overall (L1-M3) and between visits (L1-M2, M2-L2, L2-M3). P-values under 0.05 are considered significant and denoted by a star. Means, standard deviations, and regression coefficients were calculated in R. All graphs were created in R. All code used in the statistical analysis is available through Columbia University Library’s Academic Commons (URL: provided upon paper acceptance).

## Results

All maternal and fetal ultrasonic measurements, patient-specific CAD models of reproductive anatomy, and estimated combined uterus and cervix volume data are available through Columbia University Library’s Academic Commons (URL: provided upon paper acceptance).

### Maternal ultrasonic measurements

Ultrasonic measurements of maternal anatomy varied across participants, and gestational age impacted each analyzed ultrasonic measurement of maternal anatomy differently. All uterine diameters increased across gestation (fig. 3). Overall, IS-UD increased by 8.2 mm/week, AP-UD by 2.1 mm/week, and LR-UD by 6.3 mm/week (tab. 2). The periods of greatest increase differed between intrauterine diameters, with the largest slope for IS-UD between L1-M2 (10.5 mm/week) and the largest slope for AP-UD and LR-UD between M2-L2 (2.4 and 8.7 mm/week, respectively).

**Fig 3.**
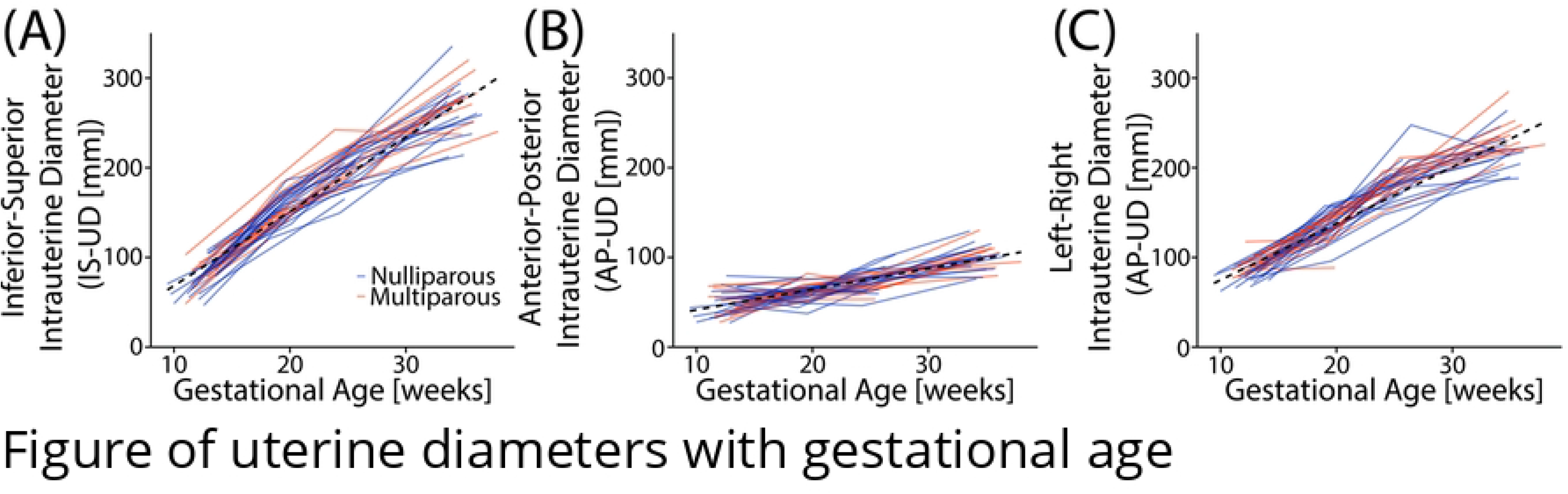
Intrauterine diameters with gestational age. (A) Inferior-superior (IS-UD), (B) anterior-posterior (AP-UD), and (C) left-right (LR-UD) intrauterine diameter with gestational age for all participants. The dotted line marks the overall slope, and participants are color-coded by parity (blue = nulliparous, orange = multiparous).

LUS-UT and CL decreased as gestational age increased. LUS-UT significantly decreased across gestation (tab. 2), with the most significant thinning between L2-M3 (−0.1 mm/week). The slope of CL decrease (fig. 4) was −0.2 mm/week (tab. 2), with the greatest and only significant decrease in L2-M3 (−0.5 cm/week). No parity-based significant differences in means or slopes were found for any maternal measurements included in the analysis (S1 Table 1). Of note, transvaginal images taken during a lower uterine segment contraction (n=18) were not included in the slope analysis.

**Fig 4.**
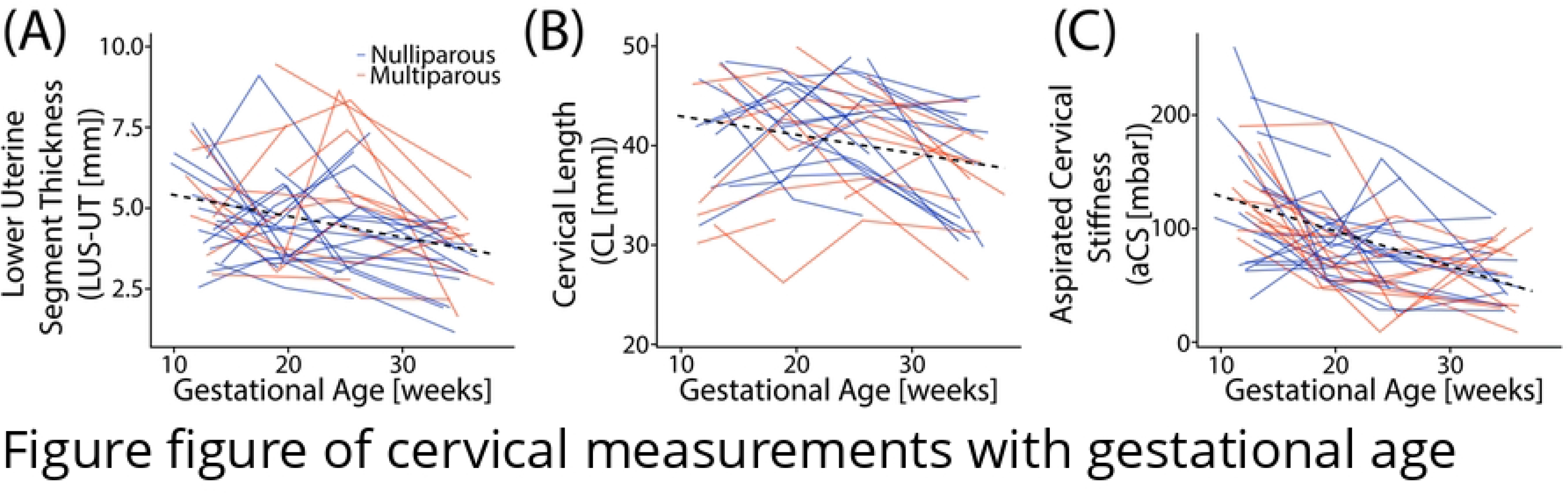
LUS-UT, CL, and aCS with gestational age. (A) Lower uterine segment thickness (LUS-UT), (B) cervical length (CL), and (C) aspirated cervical stiffness (aCS) with gestational age for all participants. The dotted line marks the overall slope, and participants are color-coded by parity (blue = nulliparous, orange = multiparous).

### Aspirated cervical stiffness measurements

The aspirated cervical stiffness (aCS) values found across participants throughout gestation fell within previously reported values for pregnancies at low-risk for PTB (fig. 4), with ranges similar to that reported in Badir et al. [18]. Across gestation, cervical stiffness decreased with increasing gestational age by −3 mbar/week (tab. 2). The only significant change in cervical stiffness between visits was L1-M2 (−5 mbar/week). This was also the largest decrease between visits. No significant differences were found in aCS value means or slopes based on parity (S1 Table 1).

### Estimated uterocervical volume and fetal weight

The estimated uterocervical volume (EUV) and fetal weight (EFW) increased for all participants across gestation (fig. 5).

**Fig 5.**
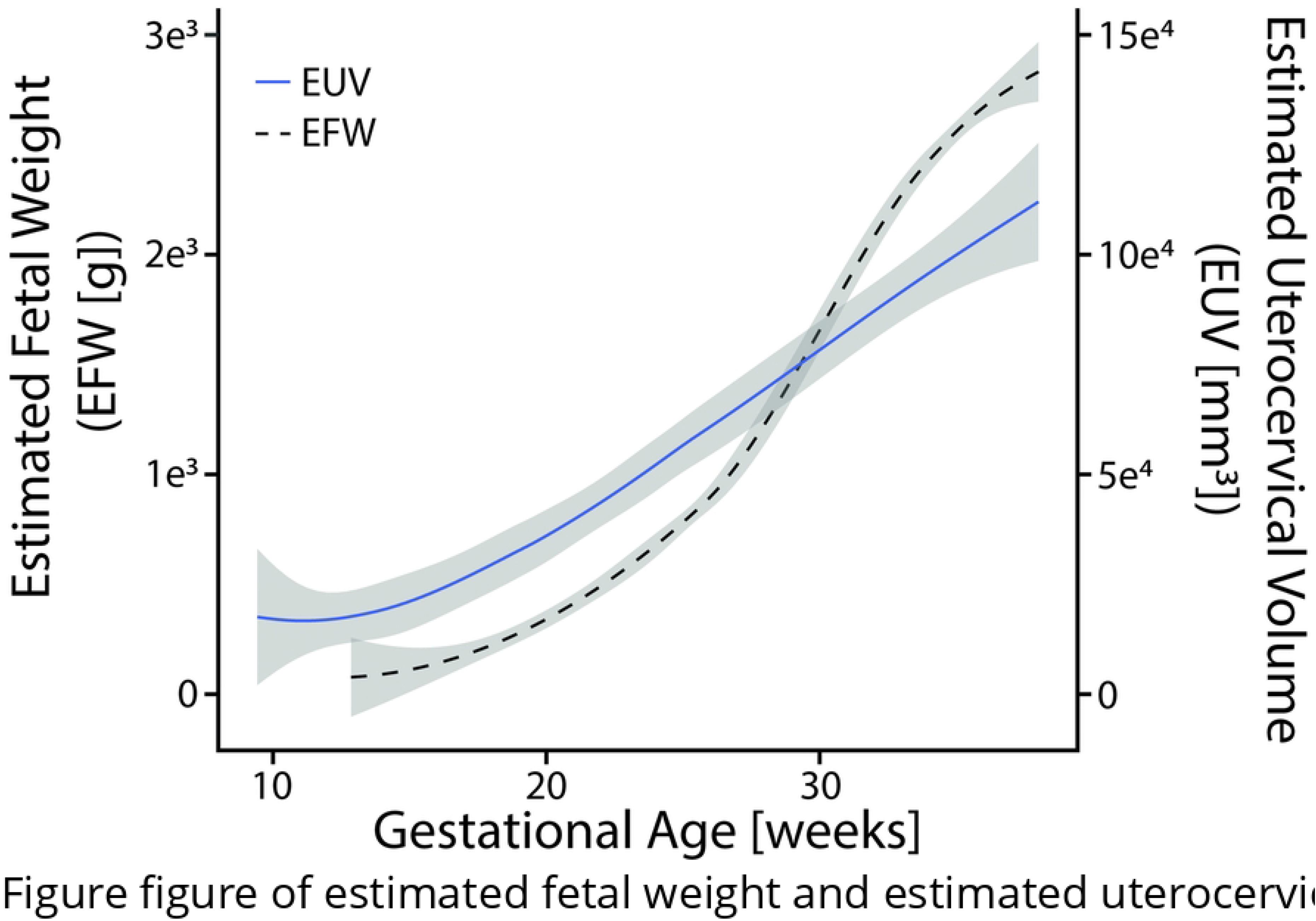
Estimated uterocervical volume (EUV) and fetal weight (EFW) across gestation. Mean estimated uterocervical volume ([mm^3^], solid blue line) and estimated fetal weight ([g], dashed black line) across gestation with standard deviation (gray).

## Discussion

This novel clinical dataset of human pregnancy presents time-course measurements of maternal uterocervical geometry, cervical stiffness, and fetal size. Participants attended four research visits from the late first to the middle third trimesters during which 2D ultrasound images were acquired of the uterus, cervix, and fetus. We found that intrauterine diameters increase with gestational age, with the rate of increase dependent on gestational age, which is logical since the uterus must grow and stretch to accommodate the growing fetus and increasing amniotic fluid volume. The largest increase was found in IS-UD, the smallest in AP-UD, and LR-UD fell in between. These findings suggest less mechanical resistance to uterine growth superior, as compared to inferior, to the uterus. This makes sense, since since the uterus is bound inferiorly by the bony pelvis while the space superior to it contains primarily soft internal organs. The mechanical rationale for why LR-UD increases more than the anterior-posterior intrauterine diameter warrants further exploration. We hypothesize this is because the AP boundaries (anterior = abdominal well, posterior = spine) are more restrictive than the lateral boundaries. In a previous study of maternal anatomy in which we investigated uterine diameters in the standing and supine positions, we found gravity-based differences in AP-UD trends. For example, the AP-UD overall, AP-UD late first to early second-trimester, and LR-UD late second to middle third slopes were significantly larger in the standing compared to the supine position (S2 Table 2) [20]. This has implications for future digital twin studies in which maternal position is considered. However, the AP-UD slope was smaller for all gestational time points than the LR-UD slope. Lastly, we found a positive relationship between all intrauterine diameters and EFW, allowing for the estimation of IS-UD, AP-UD, and LR-UD from EFW in scenarios where extended-field-of-view imaging is not available, and exact intrauterine diameter values are not necessary (fig. 6).

**Table 2.**
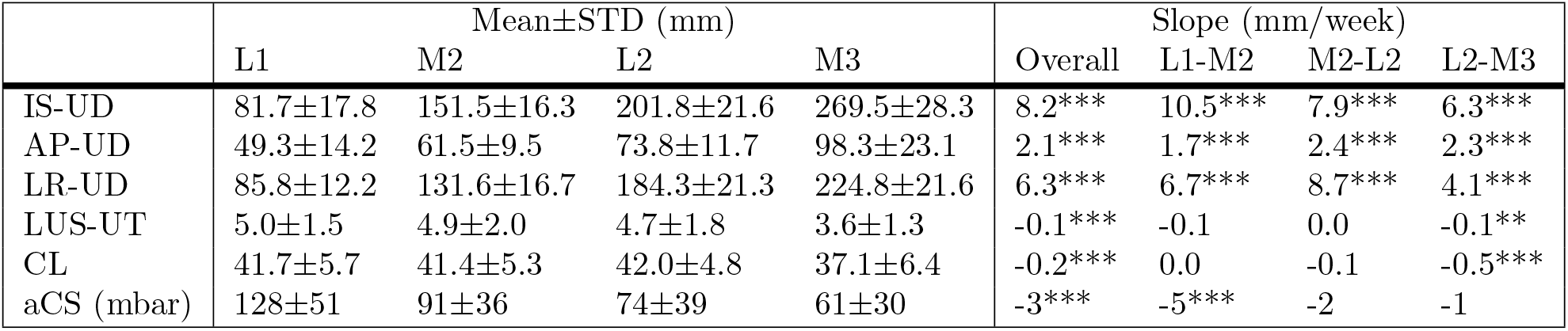
Mean dimension measurements and slopes. Mean and standard deviation (STD) for 2D ultrasonic and cervical stiffness measurements across all participants and gestation ages. Overall (L1-M3) and between visit slopes for 2D ultrasonic measurements with participant correlation. Asterisks indicate significance level: *P <*.05 ( ), *P* = .01-.05 (*), *P* = .001-.01(**), *P <*.001 (***).

|  | Mean±STD (mm) |  |  |  | Slope (mm/week) |  |  |  |
| --- | --- | --- | --- | --- | --- | --- | --- | --- |
|  | L1 | M2 | L2 | M3 | Overall | L1-M2 | M2-L2 | L2-M3 |
| IS-UD | 81.7±17.8 | 151.5±16.3 | 201.8±21.6 | 269.5±28.3 | 8.2*** | 10.5*** | 7.9*** | 6.3*** |
| AP-UD | 49.3±14.2 | 61.5±9.5 | 73.8±11.7 | 98.3±23.1 | 2.1*** | 1.7*** | 2.4*** | 2.3*** |
| LR-UD | 85.8±12.2 | 131.6±16.7 | 184.3±21.3 | 224.8±21.6 | 6.3*** | 6.7*** | 8.7*** | 4.1*** |
| LUS-UT | 5.0±1.5 | 4.9±2.0 | 4.7±1.8 | 3.6±1.3 | -0.1*** | -0.1 | 0.0 | -0.1** |
| CL | 41.7±5.7 | 41.4±5.3 | 42.0±4.8 | 37.1±6.4 | -0.2*** | 0.0 | -0.1 | -0.5*** |
| aCS (mbar) | 128±51 | 91±36 | 74±39 | 61±30 | -3*** | -5*** | -2 | -1 |

**Fig 6.**
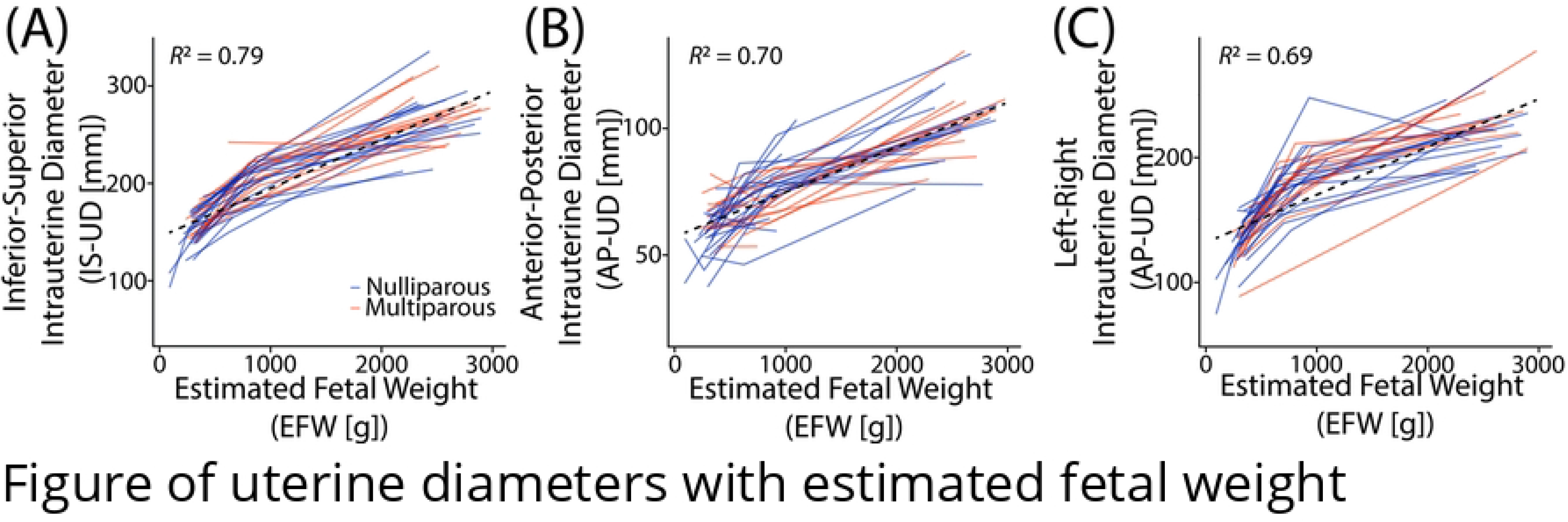
Intrauterine diameters graphed with estimated fetal weight. (A) Inferior-superior (IS-UD), (B) anterior-posterior (AP-UD), and (C) left-right (LR-UD) intrauterine diameters with corresponding estimated fetal weight (EFW). A linear regression was fit for each, with R^2^ values between 0.69 − 0.79 for all. Individuals were plotted according to parity (orange = nulliparous, blue = multiparous).

Features of cervical remodeling palpable to the clinician include decreasing cervical stiffness and length. Unsurprisingly, we found that aCS decreased with CL and LUS-UT. For example, the LUS-UT decrease was statistically significant across gestation and between the late second and middle third trimesters (tab. 2). The aCS and CL decreased significantly across gestation, though not at the same rate. The cervix softens the most between the late first and middle second trimester, whereas the cervix shortens the most between the late second and early third trimester (tab. 2). This trend of cervical softening before shortening has been reported in previous cervical aspiration studies in humans [18, 28]. Numerous studies in rodents and macaques support this finding and provide insight into the underlying tissue changes [29, 30]. Specifically, though the collagen content (per dry weight) in the cervix remains constant during pregnancy, its hierarchical structure remodels drastically through constant synthesis, assembly, and degradation of collagen fiber components [30–32]; in the nonpregnant cervix, collagen fibers are highly organized and cross-linked. As pregnancy progresses, collagen crosslink maturity declines the most in mid-pregnancy, corresponding to the fastest decline in tissue stiffness [30, 32]. Our work demonstrating the dynamic profile of cervical length and stiffness changes suggests that, while the cervix softens early in pregnancy, the mechanical load from the uterine wall’s pull and the amniotic sac’s push is not large enough to deform it until the late second trimester when the LUS thin and cervix shortens at its fastest rate as fetal growth accelerations (fig. 5).

### Comparisons to previously published data

A detailed comparison between the data presented and an existing longitudinal cohort study is presented in S2 Appendix [20]. Both cohorts demonstrate a significant difference in overall LUS-UT slope, with the existing cohort exhibiting a statistically significant decrease between each research visit [20]. However, no significant differences for between visit slopes are found between cohorts, further supporting our finding that LUS-UT thins most beginning the late second trimester (tab. S2 Table 3).

Several other existing studies involving measurements of lower uterine segment wall thickness across gestation agree with our findings. Degani et al. reported a significant decrease in LUS thickness in 25 uncomplicated pregnancies at 8-week intervals between 15-39 weeks gestation [33]. Similarly, Ginsberg et al. found the lower uterine segment to thin with increasing gestational age in a cross-sectional study of 350 singleton uncomplicated pregnancies from 15 to 42 weeks gestation [34]. Neither of these studies analyzed rate of thinning [33, 34]. Durnwald et al. evaluated LUS in a cross-sectional study of 175 singleton uncomplicated pregnancies in the first, second, and third trimesters [35]. Dissimilar to our findings, they reported significant lower uterine segment wall thinning between the first and second, but not between the second and third, trimester [35]. A key methodological difference between Durnwald et al. and us is performance of the LUS measurement 2cm above the internal os in a TA image with a full bladder, whereas we measured the thinnest segment of the anterior uterine wall in a TV image with an empty bladder [35].

Many previous studies show that the cervix softens during pregnancy. Using the same linear regression approach on an existing dataset of shear wave speed across pregnancy by Carlson et al., the overall slope decreased across gestation, with the largest decrease occurring between the late first and early second trimesters, concurring with the aCS slopes presented here (S2 Table 3) [21]. Carlson et al. also found that the greatest decrease in shear wave speed occurred closest to the proximal cervix, with a significant non-linear trend, and no significant trends in the distal cervix [21].

Longitudinal elastography studies have shown softening throughout pregnancy [19, 36–40], though no others noted significant early softening. It seems important to establish a normal rate of softening, especially early in pregnancy, because premature and/or accelerated softening is logically associated with risk of PTB. For example, a cervical aspiration study demonstrated that patients presenting for cerclage due to a history of preterm birth and short cervical length have significantly softer cervices than normal controls [28]. Another demonstration of softening early in normal pregnancy is provided by Badir et al. in a study of 100 patients at low risk for PTB [18]. Importantly, the aCS mean and standard deviation reported in that study overlat with that of ours (S2 Fig. 1). In summary, the agreement across studies of progression in cervical softening is promising and warrants further material modeling efforts to determine relationships in cervical stiffness changes measured via different modalities.

### Limitations

Though this work presents a novel dataset of maternal and fetal growth with changes to cervical stiffness across pregnancy, it has limitations. Our participants were predominantly white, thus study of more diverse populations is imperative. Further, the dataset is incomplete, with missed participant visits due to scheduling conflicts, inclement weather, participant dropouts due to changing providers, the coronavirus pandemic, and being lost to follow-up. Scheduling conflicts also led to lack of cervical aspiration measurements at some research visits due to unavailability of trained physicians. Further, while the aCS differences were observed in some patients across the three average measurements, with ranges greater than 50 mbar for 16 L1 measurements, 3 M2 measurements, 1 L2 measurement, and 1 M3 measurement are likely due to the viscoelastic behavior of the cervix, it is possible that other factors could contribute to these measurement ranges. These include cervical surface friction, force applied to the probe during measurement, and differences in inter-operator technique, all of which deserve focused evaluation in future studies. Ultrasonic measurements may be compromised by inter-operator variability [20], though we attempted to minimize this by having an experience clinician verify all dimensions. In some of the research visits, the clinician deemed dimensions “unmeasurable” due to poor visualization and/or acquisition. While this reflects the “real world” situation, these measurements were excluded from statistical analysis, though estimates were used to generate the solid models for the estimated uterocervical volume and are included in the available dataset (data sets published by Columbia Academic commons; URL provided up acceptance). Finally, measurements of estimated uterocervical volume have not been verified for gestational ages of less than 37 weeks, and thus it is unknown how errors in ultrasonic measurements may propagate to errors in estimated uterocervical volume.

## Conclusion

This work presents holistic measurements of changes to maternal anatomy with progressive growth of the reproductive structures and fetus during pregnancy on a patient-specific basis, with complementary measurements of cervical softening. Anatomic measurements of the uterus, cervix, and fetus were collected using typical clinical B-mode 2D ultrasound acquisitions. The uterine diameter increased most in the inferior-superior direction and least in the anterior-posterior direction. The lower uterine segment thinned significantly across gestation as the cervical stiffness and length decreased. The most rapid cervical softening occurred between the late first and middle second trimester, whereas the greatest shortening occurred between the late second and middle third trimester. These findings quantitatively characterize changes in the uterus and cervix in normal pregnancy, providing a foundation for *in-silico* studies of digital twins that can be manipulated to model multiple pregnancy situations.

## Supporting information

### S1 Appendix. Supplemental Methods

In addition to the methods outlined in this work, the authors provide further details on the available data, Columbia University (CU) and Intermountain Healthcare (IH) cohorts, and estimation of uterocervical volume from patient-specific computer-aided design models (digital twins). The IH cohort was originally published in [20].

Not all participants in the Columbia University cohort could make it to all four research visits, and not all research visits could include cervical stiffness measurement. A summary of how many participants attended each research visit timeframe and the number of ultrasonic and aspiration measurements collected is presented in S1 Table 1. The number who attended the prior visit is also given, which is pertinent for the between-visit analysis.

**S1 Table 1.** Research visit attendance. Summary of the number of participants who attended during each visit timeframe and the number who attended the previous visit.

|  | L1 (9<sup>3</sup>-15<sup>3</sup>) | M2 (17<sup>3</sup>-20<sup>5</sup>) | L2 (23<sup>3</sup>-27<sup>1</sup>) | M3 (33<sup>4</sup>-38<sup>0</sup>) |
| --- | --- | --- | --- | --- |
| **Attended visit** | 45 | 37 | 34 | 29 |
| Ultrasound collected | 45 | 37 | 33 | 28 |
| Aspiration collected | 43 | 35 | 29 | 24 |
| **Attended prior visit** | N/A | 35 | 32 | 28 |

Though the between-visit slope in the CU and IH cohorts were directly compared, the gestational age range for each visit did not exactly overlap. In the case of visit 2, there is no overlap in gestational age between the two cohorts. The gestational age range and number of participants for each visit in the CU and IH cohorts are given in S2 Table 2.

**S1 Table 2.** Visit timeframes for Columbia University (CU) and Intermountain Healthcare (IH) cohorts. Some overlap in gestational age exists for all research visits, with the exception of visit 2.

|  | Visit 1 | Visit 2 | Visit 3 | Visit 4 |
| --- | --- | --- | --- | --- |
| CU Gestational Age | L1 (9<sup>3</sup>-15<sup>3</sup>) | M2 (17<sup>3</sup>-20<sup>5</sup>) | L2 (23<sup>3</sup>-27<sup>1</sup>) | M3 (33<sup>4</sup>-38<sup>0</sup>) |
| CU participants | 45 | 37 | 34 | 28 |
| IH Gestational Age | L1 (8<sup>3</sup>-14<sup>5</sup>) | E2 (15<sup>3</sup>-16<sup>6</sup>) | L2 (22<sup>0</sup>-24<sup>6</sup>) | M3 (32<sup>0</sup>-34<sup>4</sup>) |
| IH participants | 29 | 29 | 29 | 29 |

The set of maternal anatomic dimensions on which linear regressions were performed was analyzed because they previously demonstrated excellent agreement between observers, except cervical length (good agreement), and represent gross uterine and cervical size. However, they do not characterize the shape of the uterus and cervix, which is necessary to generate parametric patient-specific computer-aided design (CAD) models. Here, we list all maternal anatomic dimensions collected and made available through Columbia University’s Academic Commons (URL provided up acceptance).

These dimensions are based on previous definitions, and all measured dimensions were checked by a maternal-fetal medicine clinician (M.H.) [20, 26]. For clarity, the set of measurements on which linear regressions were performed were referred to using anatomic descriptors, i.e. inferior-superior intrauterine diameter is IS-UD. Previous work did not use this naming approach, so we present the anatomic descriptors in bold next to the equivalent dimensions.

From the transabdominal sagittal ultrasonic image, the following dimensions were measured:

- UD1 **(IS-UD)**: longest inferior-superior intrauterine diameter was measured from the fundal to the lower uterine segment endometrium (S1 Fig. 1a)
- UD2: anterior intrauterine diameter, measured perpendicularly from midpoint of UD1 to anterior intrauterine wall (S1 Fig. 1a)
- UD3: posterior intrauterine diameter, measured perpendicularly from midpoint of UD1 to posterior intrauterine wall (S1 Fig. 1a)
- UD23 **(AP-UD)**: sum of UD2 and UD3 (S1 Fig. 1a)
- UT1: fundal uterine wall thickness, measured as close to the superior end of UD1 as possible (S1 Fig. 1a)
- UT2: anterior uterine wall thickness, measured at the anterior end of UD2 (S1 Fig. 1a)
- PCO: perpendicular cervical offset, perpendicular distance between UD1 and internal os (S1 Fig. 1a)
- UD3a eq: posterior dimension perpendicular to UD1 at 25% of UD1 from the superior intrauterine wall (S1 Fig. 1a)
- UD3b eq: posterior dimension perpendicular to UD1 at 75% of UD1 from the superior intrauterine wall (S1 Fig. 1a)
- UD3a ex: extrema perpendicular distance between posterior wall and UD1 superior to UD3, no always applicable (S1 Fig. 1b)
- UD1a ex: distance from inferior end of UD1 to UD3a ex, only measured if UD3a ex was measured (S1 Fig. 1b)
- UD3b ex: extrema perpendicular distance between posterior wall and UD1 inferior to UD3, no always applicable (S1 Fig. 1b)
- UD1a ex: distance from inferior end of UD1 to UD3b ex, only measured if UD3b ex was measured (S1 Fig. 1b)

**S1 Fig. 1 Solid Modeling Measurements**. (A) Measurements collected from sagittal ultrasound images to capture the sagittal uterine shape and placement of the cervix, with measurements of posterior intrauterine diameter taken equidistantly (eq) along UD1. (B) Alternative method to collecting posterior intrauterine diameter measurements, taken at the superior (UD3a) and inferior (UD3b) extrema (ex). (C) Measurements collected from axial ultrasound images to capture the axial uterine shape. (D) Measurements taken from sagittal transvaginal ultrasounds to capture the uterus and cervix. (E) Measurements of outer cervical diameter taken equidistantly along the cervical length. (F) Measurements of inner cervical diameter taken equidistantly along the cervical length.

From the TA axial ultrasonic image, the following dimensions were measured:

- UD4 **(LR-UD)**: longest left-right intrauterine diameter (S1 Fig. 1c)
- UT3: left/right uterine wall thickness, measured as close to the left or right end of UD4 as possible (S1 Fig. 1c)

From the TV sagittal image, the following dimensions were measured:

- UT4 **(LUS-UT)**: lower uterine segment thickness, measured as the thinnest portion of the visible anterior uterine wall (S1 Fig. 1d)
- **CL**: cervical length, measured as the distance between the internal os (where the anterior and posterior cervix meet in the image) and the external os (S1 Fig. 1d)
- AUCA: anterior uterocervical angle, measured as the angle between the lower uterine segment and cervical canal, placed as 1 cm lines starting at the internal os along the anterior uterine wall and the cervical canal (S1 Fig. 1d)
- CD1 25: outer cervical diameter measured at 25% of CL from the internal os (S1 Fig. 1e)
- CD1 50: outer cervical diameter measured at 50% of CL from the internal os (S1 Fig. 1e)
- CD1 75: outer cervical diameter measured at 75% of CL from the internal os (S1 Fig. 1e)
- CD2 25: inner cervical diameter measured at 25% of CL from the internal os (S1 Fig. 1f)
- CD2 50: inner cervical diameter measured at 50% of CL from the internal os (S1 Fig. 1f)
- CD2 75: inner cervical diameter measured at 75% of CL from the internal os (S1 Fig. 1f)

Of these maternal anatomic dimensions, the only ones not used in the parametric patient-specific models are CD1 25, CD1 75, CD2 25, and CD2 75. Future iterations of the modeling approach may include them to attain a more refined cervical shape.

The approach to modeling the uterus was updated slightly from the previous method [20]. First, due to the variability in sagittal posterior uterine wall shape observed between patients and gestational ages, several methods of building the posterior wall were generated based on parametric measurements collected (updating fig.5a in [20]). If both the superior (UD3a) and inferior (UD3b) posterior diameter measurements were collected as extremum (ex), quarter ellipses were used at the inferior and superior ends of the uterus with a spline connecting them through the middle posterior diameter (UD3) (S1 Fig. 2a). A spline was used for the entire posterior uterine profile if no posterior diameters were measured as extremum, thus using the equidistant (eq) approach (S1 Fig. 2b). Finally, if only one of the inferior (S1 Fig. 2c) or superior (S1 Fig. 2d) posterior diameter measurements were collected as extremum, a quarter ellipse was used at the end with the extremum measured, and a spline connecting the middle posterior diameter and equally placed measurement of posterior diameter to the end of the inferior-superior axis.

**S1 Fig. 2 Sagittal uterine modeling approaches**. Method to model the sagittal uterus when (A) both superior (UD3a ex) and inferior (UD3b ex) posterior intrauterine diameters are taken as extrema, (B) both superior (UD3a eq) and inferior (UD3b b) are taken equidistantly along the inferior-superior axis, (C) only the inferior (UD3b ex) is taken as extrema, and (D) only the superior (UD3a ex) is taken as extrema. Red arrows depict the ends of the spline, where tangency constraints are enforced.

The method of generating the uterine body was also updated, with an inferior-superior loft through axial profiles used rather than a left-right loft function, updating fig.5c in [20]. The coronal uterine shape was still modeled as an ellipse, with elliptical profiles placed coincidentally with UD3a and UD3b (**??**a). The intrauterine diameters at these locations were assigned by their location within the coronal ellipse defined by UD1 and UD4, with UD4a given by eq. 1 and UD4b given by eq. 2.

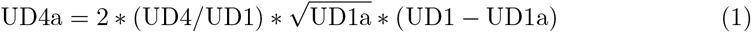

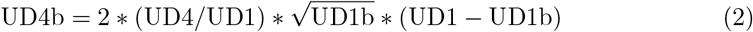

For the outer uterus, eq. 1 and eq. 2 were modified such that the superior and inferior uterus were defined to include uterine wall thicknesses (UT1 and UT4, respectively). The anterior-posterior diameter of the axial elliptical profiles was defined by the sagittal uterine profile (**??**b), and the loft function executed using the sagittal profile as guides (**??**c).

**S1 Fig. 3 Uterine body modeling approach**. The uterine body modeling approach was updated to enforce an elliptical axial cross section. (A) Left-right diameters were placed coincidentally at locations with a posterior intrauterine diameter to generate an elliptical coronal cross section. (B) Axial ellipses are generated at each location with a posterior intrauterine diameter to be defined by the left-right intrauterine diameters and sagittal uterine profile. (C) The uterine body is generated using the loft functions, with the sagittal shape acting as guides and the axial ellipses acting as loft profiles.

### S2 Appendix. Additional Analyses

In addition to the analyses on changes in intrauterine diameter, lower uterine segment thickness, and cervical length with gestational age, analyses on the effects of parity and maternal position were performed. Comparison between cohorts and to existing datasets was also undertaken to assess our findings.

All measurements of fetal anatomy and amniotic fluid levels were compared to clinically established growth charts (data sets published by Columbia Academic commons; URL provided up acceptance), and few anomalies were identified. Crown-rump length measurements were compared against the INTERGROWTH-21st Project’s crown-rump length percentiles during the first trimester, with 40 out of 43 measurements within the 5th-95th percentile [41]. Similarly, 95 out of 101 measurements of estimated fetal weight were within the 5th-95th percentile based on data from the World Health Organization [42]. Finally, 93 out of 100 maximum vertical pocket measurements and 52 out of 57 amniotic fluid index measurements were within the 5th-95th percentile based on data from the National Institute of Child Health and Development [43]. Thus, the pregnancies analyzed in the study are, for the most part, within normal ranges.

In analyzing parity in the Columbia University (CU) cohort, presented in this study, no significant difference was found in mean for any of the analyzed measurements at any gestational age between nulliparous and multiparous participants (S1 Table 1). This aligns with previous work, which reports no significant difference in lower uterine segment thickness based on parity [33, 34]. However, it is contradictory to the findings of Durnwald et al., who reported a significant difference between nulliparous and multiparous lower uterine segment thicknesses, though these results appear to include measurements from all three trimesters [35].

**S2 Table 1.** Effect of parity on maternal ultrasonic measurements. Mean ± standard deviation for nulliparous and multiparous participants during the late first (L1), middle second (M2), late second (L2), and middle third (M3) trimester with the corresponding Welch’s T-test p-value. Performed for inferior-superior (IS-UD), anterior-posterior (AP-UD), and left-right (LR-UD) intrauterine diameters and lower uterine segment thickness (LUS-UT), cervical length (CL), and aspirated cervical stiffness (aCS).

|  | IS-UD | AP-UD | LR-UD | LUS-UT | CL | aCS |
| --- | --- | --- | --- | --- | --- | --- |
| L1 Mean ± STD (mm) | | | | | | |
| Multiparous | 83.6±15.7 | 45.2±14.6 | 88.9±10.8 | 7.8±4.0 | 47.5±11.4 | 131±34 |
| Nulliparous | 80.6±19.1 | 50.1±14.1 | 83.9±12.8 | 7.5±7.2 | 47.4±8.8 | 127±61 |
| Parity T-test | 0.56 | 0.27 | 0.17 | 0.88 | 0.99 | 0.77 |
| E2 Mean ± STD (mm) | | | | | | |
| Multiparous | 151.4±16.4 | 65.1±8.6 | 129.2±18.2 | 5.8±3.0 | 44.8±11.1 | 87±40 |
| Nulliparous | 151.5±16.6 | 59.5±9.6 | 132.9±16.1 | 6.4±4.6 | 44.5±5.7 | 94±34 |
| Parity T-test | 0.98 | 0.08 | 0.54 | 0.61 | 0.91 | 0.58 |
| L2 Mean ± STD (mm) | | | | | | |
| Multiparous | 208.0±19.3 | 70.5±9.9 | 187.1±17.4 | 5.8±2.7 | 42.4±7.3 | 59±29 |
| Nulliparous | 198.6±22.4 | 75.8±12.4 | 182.8±23.2 | 5.2±4.2 | 45.1±6.8 | 83±41 |
| Parity T-test | 0.23 | 0.19 | 0.56 | 0.58 | 0.31 | 0.08 |
| M3 Mean ± STD (mm) | | | | | | |
| Multiparous | 275.8±24.7 | 103.6±12.9 | 232.8±21.9 | 3.9±1.4 | 36.0±7.0 | 61±32 |
| Nulliparous | 264.7±30.6 | 100.5±14.3 | 218.9±20.1 | 3.4±1.1 | 38.7±6.5 | 60±28 |
| Parity T-test | 0.30 | 0.55 | 0.10 | 0.27 | 0.32 | 0.98 |

Given the difference between visit slopes found in the CU cohort, the same analysis was undertaken in the Intermountain Healthcare (IH) cohort to compare slopes in the standing and supine position (S2 Table 2). Previously, it was reported that position significantly affected slopes for all intrauterine diameters, but not lower uterine segment thickness or cervical length [20]. The patient-correlated linear regression analysis approach found no significant difference in the inferior-superior intrauterine diameter (IS-UD) overall or between any visits. This approach found a significant difference in the overall slope of anterior-posterior intrauterine diameter (AP-UD) and L1-E2 slope, with the supine slope being smaller than the standing slope in both cases. The only significant difference found in left-right intrauterine diameter slope (LR-UD) was found between L2-M3, with the supine slope being smaller than the standing slope. No significant difference was found in lower uterine segment thickness (LUS-UT) and combined cervical and isthmus length (CL+IS) for any slope between standing and supine.

**S2 Table 2.** Effect of position on overall and between visit slope. Overall and between visit slopes for Intermountain Healthcare (IH) cohort 2D ultrasonic measurements with participant correlation in the standing and supine position with Welch’s T-test p-value. Visit occurred in the late first (L1), early second (E2), late second (L2), and middle third (M3) trimesters. Asterisks indicate significance level compared to the null hypothesis (slope is zero): *P <*.05 ( ), *P* = .01-.05 (*), *P* = .001-.01(**), *P <*.001 (***).

|  | Overall Slope (mm/week) |  |  | L1-E2 Slope (mm/week) |  |  | E2-L2 Slope (mm/week) |  |  | L2-M3 Slope (mm/week) |  |  |
| --- | --- | --- | --- | --- | --- | --- | --- | --- | --- | --- | --- | --- |
|  | Supine | Standing | p-val | Supine | Standing | p-val | Supine | Standing | p-val | Supine | Standing | p-val |
| IS-UD | 8.7*** | 9.0*** | 0.82 | 7.9*** | 7.8*** | 0.26 | 9.1*** | 9.2*** | 0.73 | 8.8*** | 9.4*** | 0.25 |
| AP-UD | 2.6*** | 3.0*** | <0.001 | 2.1*** | 4.3*** | <0.001 | 2.6*** | 2.2*** | 0.39 | 2.7*** | 2.9*** | 0.56 |
| LR-UD | 7.6*** | 8.2*** | 0.57 | 5.8*** | 5.5*** | 0.74 | 9.5*** | 8.4*** | 0.21 | 6.8*** | 9.2*** | 0.008 |
| LUS-UT | -0.32*** | -0.35*** | 0.94 | -0.7* | -0.5 | 0.65 | -0.5* | -0.5* | 0.78 | -0.2* | -0.2* | 0.46 |
| CL+IS | -0.5*** | -0.6*** | 0.87 | 0.1 | -0.3 | 0.74 | -0.5 | -0.6 | 0.85 | -0.9** | -0.9** | 0.95 |

The changes in maternal ultrasonic measurements were compared to the existing serial dataset collected at Intermountain Healthcare (IH) in Provo, Utah [20]. Similar to this study, patients at low risk for PTB underwent the same ultrasonic scanning protocol in the late first, early second, late second, and middle third trimesters, with images collected with participants in both the standing and supine position. Cervical stiffness was measured at each visit via ultrasonic shear wave speed (SWS) [21]. The original analysis of these data presented linear mixed effects models for each ultrasonic parameter to estimate its relationship with gestational age, position, and parity [20]. To directly compare the presented dataset (CU) and previous dataset (IH), the same patient-correlated linear regression analysis was performed, and a Welch’s t-test was done to find significant differences (tab. S2 Table 3). Cervical length was combined with isthmus length (CL+IS) in the IH cohort to match the CU study protocol. Also, all CU cervical and lower uterine segment measurements were included for comparison, as the presence of contraction in TV images is considered a subjective finding and methods for identification may not be the same between cohorts [44].

In Carlson et al., shear wave speed (SWS) measurements were collected at 5 gestational ages to measure changes in cervical stiffness [21]. The first four visits coincide with those from Louwagie et al., thus the fifth visit was not included in this analysis [20]. At each visit, three SWS images were taken, and split into 4 regions in the anterior cervix: proximal (P), mid-proximal 1 (MP1), mid-proximal 2 (MP2), and middle (M) [21]. All SWS data is publicly available, and each image produces 100-200 SWS values in each of the four cervical regions. To determine a shear wave speed measurement for each participant at each visit, all P, MP1, MP2, and M measurements were averaged across all values from all 3 SWS images. To quantify the relationship between gestational age and SWS, 4 linear regressions were performed to find the overall slope and slope between visits 1-2, 2-3, and 3-4. The data averaging and the linear regressions were both performed in RStudio version 2023.6.0.421 [27].

**S2 Table 3.** Comparison between Columbia University (CU) and Intermountain Healthcare (IH) cohorts [20]. Overall and between visit slopes for IH cohort in supine position, with Welch’s t-test *P* -values against CU cohort (*P* -value).

|  | Overall (mm/week) |  | L1-E2 (mm/week) |  | E2-L2 (mm/week) |  | L2-M3 (mm/week) |  |
| --- | --- | --- | --- | --- | --- | --- | --- | --- |
|  | IH Slope | *P*-value | IH Slope | *P*-value | IH Slope | *P*-value | IH Slope | *P*-value |
| IS-UD | 8.7*** | 0.75 | 7.9*** | 0.03 | 9.1*** | 0.79 | 8.8*** | 0.18 |
| AP-UD | 2.6*** | <.001 | 2.1*** | 0.67 | 2.6*** | 0.15 | 2.7*** | 0.62 |
| LR-UD | 7.6*** | 0.89 | 5.8*** | 0.29 | 9.5*** | 0.31 | 6.8*** | <.001 |
| LUS-UT | -0.3*** | <.001 | -0.7* | 0.58 | -0.5* | 0.23 | -0.2* | 0.67 |
| CL+IS | -0.5*** | 0.24 | 0.11 | 0.03 | -0.5 | 0.57 | -0.9** | 0.31 |
| SWS (m/s) | -0.07*** | N/A | -0.13 | N/A | -0.10* | N/A | -0.05* | N/A |

Overall slopes were significantly different for AP-UD and LUS-UT, though the qualitative findings remain the same: AP-UD increases across gestation, and LUS-UT decreases (S2 Table 3). Additional differences are observed in the timeframe of the greatest increase in intrauterine diameters, with IS-UD increasing most between E2-L2, AP-UD increasing most from L2-M3, and LR-UD increasing most from E2-L2. That said, significant differences were only found for IS-UD between L1-E2 timeframe and LR-UD between L2-M3. A notable difference is also observed in LUS-UT slopes across gestational age, with significant decreases observed between each research visit, the largest of which occurs between L1-E2. However, no significant difference between the CU and IH datasets for LUS-UT between visit slopes was found.

CL significantly decreased between L2-M3 in the IH cohort, with no significant difference to the CU cohort. Differences in LUS-UT definition may also contribute to the inconsistencies between datasets, where LUS-UT was specifically measured in the thinnest portion of the lower uterine segment in the CU cohort but not in the IH cohort. Cervical stiffness was also collected in the IH cohort, but since SWS was measured and not aspiration, no direct quantitative comparisons can be made without a non-linear, time-dependent material model. However, the overall SWS slope decreased across gestation, with the largest (though insignificant) decrease occurring between L1-E2, which concur with the CU cohort’s aCS slopes (S2 Table 3).

Changes in cervical stiffness with gestation found in the CU cohort were compared to previous findings in aspirated cervical stiffness (aCS) and shear wave speed (SWS) [18, 21]. No raw data was publicly available for aCS with gestational age, so the published figure on mean aCS values with gestational age was reproduced with the CU cohort overlaid (S2 Fig. 1). Plotting the aCS data as such, there appears to be good overlap between the two datasets, with the means of both falling within one standard deviation of the closest gestational age.

**S2 Fig. 1 Aspirated cervical stiffness for CU cohort and Badir et al. 2013**. Mean and standard deviation of aspirated stiffness value across gestation in the Columbia University (CU) cohort and data from Badir et al. [18].

## Acknowledgments

The authors thank all research participants for their involvement in this study. The authors also thank Ivette Miranda, Ilona Karalnik, and Vanessa Pichardo for ultrasound image collection and Jordana Graifman, Michelle Vanegas, and Luiza Kalemi for research coordination. Research reported in this publication was supported by the Eunice Kennedy Shriver National Institute Of Child Health & Human Development under Award Number R01HD091153 to KMM. The content is solely the responsibility of the authors and does not necessarily represent the official views of the National Institutes of Health.

## Notes

### Competing Interest Statement

The authors have declared no competing interest.

